# Propagation electrodynamics and differential conduction of action potentials in geometrically branched squid giant axons and neurons

**DOI:** 10.64898/2026.08.03.742547

**Authors:** Xi Liu, Wenxi Fang, Ken Perlin

## Abstract

Classical neuronal cable theory relies on quasi-static electric field approximations and neglects magnetic induction, Lorentz force coupling, and transient electromagnetic currents, limiting its ability to fully characterize action potential propagation within geometrically branched axons and dendrites. This work develops a coupled Maxwell-electromagnetic cable framework by integrating finite-difference time-domain (FDTD) solutions of Maxwell’s equations with extended Hodgkin-Huxley and Fitzhugh-Nagumo membrane dynamics, incorporating magnetic gating perturbations, electromagnetic trans-membrane currents *I*_EM_, and nanoscale quantum corrections for thin neural segments. Controlled propagation experiments are designed to quantify deviations from standard cable predictions across asymmetric and symmetric axonal bifurcation geometries. Numerical results demonstrate that inductive magnetic effects lower the critical branch radius for junction conduction failure and break symmetric action potential invasion in geometrically identical child branches under external transverse magnetic fields. An electromagnetic corrected geometric ratio *GR*_EM_ is proposed to revise impedance-matching conditions at branch points, accounting for size-dependent axial current imbalance induced by magnetic and displacement currents. Parent axon conduction velocity deviates substantially from the canonical 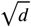 scaling law when electromagnetic feedback and quantum charge distributions are included, triggering early signal blockage at large cable diameters. Collectively, this study establishes that quasi-static cable models underestimate electromagnetic corrections to propagation speed, waveform shape, and bifurcation transmission fidelity; the coupled Maxwell-cable framework provides a comprehensive multi-physics tool for modeling electrodynamic signal behavior in complex neuronal architectures.

## 1. Introduction

The interneuron communication and data processing in the brain depends on the signal propagation among cells that involves varying geometry, which can be modeled by cable theory [1]. Interneuron communication includes electrical and chemical synapses. Electrical synapses involve direct connections between the presynaptic and post-synaptic cell membranes via gap junctions that allow the flow of electric current between cells, enabling rapid signal transmission and action potential propagation [2]. In a chemical synapse, the electrical activity of the presynaptic neuron triggers the release of neurotransmitters such as glutamate, *γ*-aminobutyric acid, acetylcholine, or norepinephrine, which bind to receptors on the postsynaptic cell (figure 1b). Computational investigations in action potentials often involves simplifying assumptions on space clamping conditions that halt the action potential propagation, uniform properties of the cable, and constant velocity propagation. These assumption can simplify the partial differential equation of the spread of membrane potential, but they cannot be used here since changing the geometry of the cable conductor affects the action potential propagation, its shape and velocity [3], [4]. The cable equation can be modified taking into account of parameters such as charge inhomogeneities [5]. The cable equation can be augmented with external driving forces from applied electromagnetic fields, synaptic excitation, and active membrane properties. The Fitzbugh-Nagumo model contains an external driving force that is dependent on potential or the spatiotemporal varying external driving force [6]. To solve the nonhomogeneous cable equation with a source or forcing term, the method of eigenfunction expansions can be used. We investigated how altering cable conductor geometry impacts action potential propagation, observing that changing the diameter of a branch can either accelerate or hinder propagation in adjacent branches, contingent on matching current flow at branch points. A geometric ratio quantifying branch diameter relationships aids in classifying equivalent cylinders, guiding analysis of propagation behavior.

**Figure 1:**
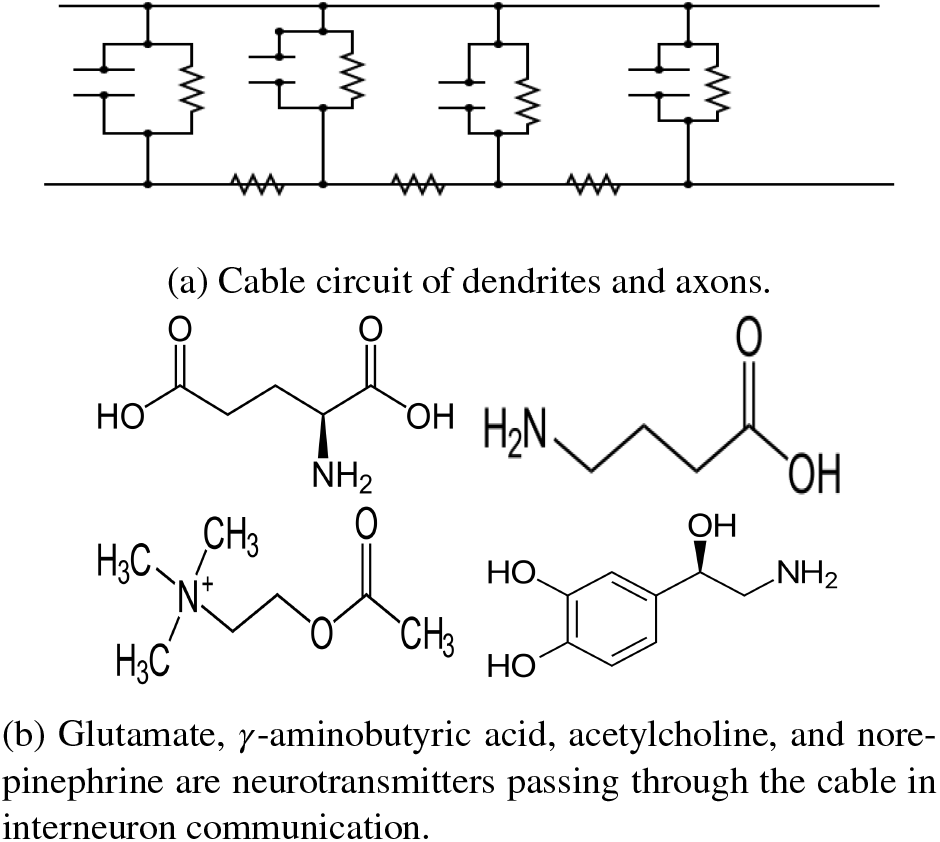
Neural cable circuit and associated neurotransmitters.

## 2. Cable theory

In cable theory, dendrites and axons are modeled by cylinders with RC circuits (containing resistors and capacitors) connected in parallel. See figure 1a for the cable circuit. The top half of the entire long cable is interfacing with the extracellular fluid, the bottom half of the entire long cable is inside cytosol, cytoplasmic matrix, or the intracellular fluid. Let *r*_*n*_ be the resistance of the *n*th RC circuit in the figure, *c*_*n*_ be the capacitance of the *n*th RC circuit, *R*_*n*_ (Ω · *cm*^2^) and *C*_*n*_ (*F* /*cm*^2^) be the specific resistance and capacitance of an unit area of membrane, *a* be the radius of the axon 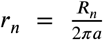, *c*_*n*_ = *C*_*n*_2*πa*. As the radius *a* of the axon increases, a greater area for current to pass through the membrane, so the resistance *r*_*n*_ becomes lower. As the circumference 2*πa* of the axon increases, more membrane can be used to store charge, so the *c*_*n*_ becomes higher. Let *ρ*_*l*_ be the specific electrical resistance of the axoplasm (cytoplasm inside the axon), then the intracellular resistance *r*_*l*_ per unit length (Ω·*cm*^−1^) in the longitudinal direction is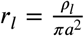. As the axon cross sectional area *πa*^2^ increases, there are more paths for the current flow the axoplasm, so the axoplasmic resistance decreases [7]. By Ohm’s law for voltage *V*, current *I*, and resistance *R, V* = *IR*

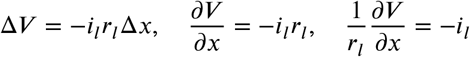

Let *i*_*n*_ be the current passing through the membrane per unit length *n*, then the total current passing through *x* units is *x* · *i*_*n*_. so the change of current in axon cytoplasm Δ*i* _*l*_ at distance Δ*x* is

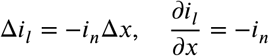

on the side of the cytoplasm, the capacitance causes a current towards the membrane, this current is displacement current 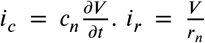 is current through the membrane. since 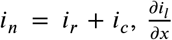 is the change of axoplasm current per unit length

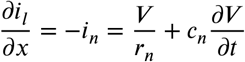

substituting 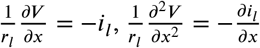

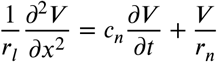

using a length constant *λ* that is a ratio of the membrane resistance *r*_*n*_ and the intracellular resistance *r*_*l*_

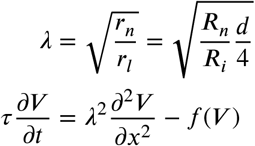

As the membrane resistance *R*_*n*_ increases, there is less current leaks across the membrane, so the space constant *λ* increases. Also, a dendrite with larger diameter *d* has a larger space constant *λ*, so the spread of current is accelerated with a larger diameter.

For the three dimensional case, let *V* be the departure of membrane potential from rest, *x* be the spatial coordinate on the core conductor, *t* be the time, *τ* be the time constant of the membrane, *λ* be a constant depending on conductor length, *f* (*x, t*) be the external driving force function. *λ* is proportional to the cable diameter 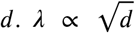. The Fitzhugh-Nagumo nerve conduction equation is [8]

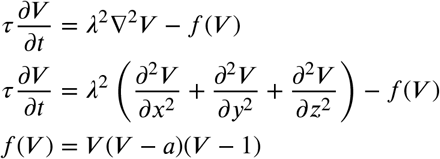

One way of solving the core conductor equation is using the eigenfunction expansion and fourier coefficients

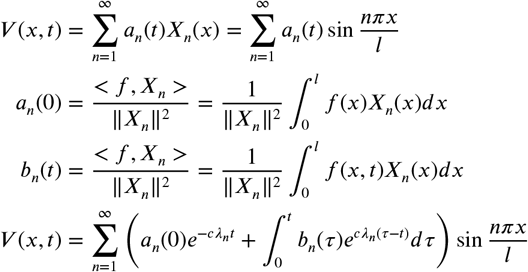

The continuity of current need to be maintained. *x*_0_ is the junction point of changing geometry. *x*_0−_ and *x*_0+_ are points immediately at the left and right of *x*_0_. *R*_*i*_ is intracellular specific resistance.

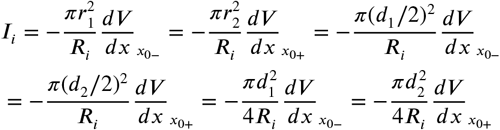

Branching usually involves a parent branch and two child branches of different radius (figure 2). To consider propagation in both directions, we distinguish that one of the branch is the starting source of the action potential propagating to the junction (usually parent branches) and the other branch in the other side of the junction (usually child branches). The propagation near the junction is having the geometric ratio

**Figure 2:**
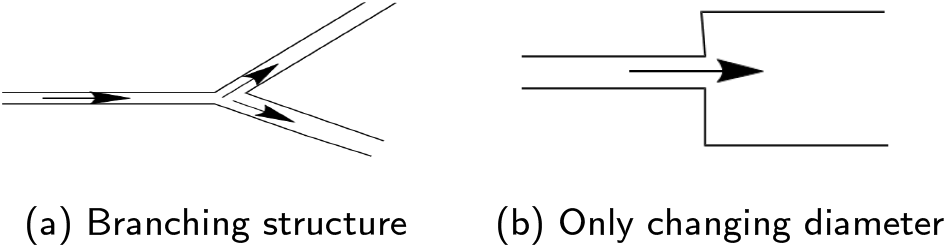
Left figure is branching, right figure is only changing diameter.

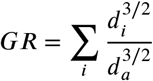

*d*_*a*_ is the diameter of the starting branch of the action potential propagating to the junction. *d*_*i*_ is the diameter of the *i*th branch on the other side of the junction. If *GR* = 1, the dendritic tree can be converted to an equivalent cylinder. If *GR <* 1, the branches collectively at the other side of the junction (usually child branches) can be converted to an equivalent cylinder with diameter smaller than the starting branch of the action potential propagating to the junction (usually parent branches). If GR > 1, the branches collectively at the other side of the junction (usually child branches) can be converted to an equivalent cylinder with diameter larger than the starting branch of the action potential propagating to the junction (usually parent branches).

## 3. Electromagnetic modified Maxwell-cable equations

Prior theoretical work has established the mathematical link between Maxwell’s electrodynamic laws and classical core-conductor cable theory, with Lindsay et al. providing a comprehensive foundational derivation mapping full electromagnetic field behavior to the standard quasistatic cable equation and outlining necessary extensions to capture inductive and displacement current terms [9, 10]. Subsequent analytical progress on coupled Maxwell-cable mixed-dimensional partial differential systems has formalised well-posedness criteria and semigroup stability analysis for radiating, geometrically curved neural cables, forming a rigorous mathematical framework for multi-physics neuronal modeling [11, 12, 13]. Modifications to the cable equation that explicitly account for transmembrane polarization and bidirectional electric field coupling were developed by Wang et al., whose work introduced field-dependent membrane current terms analogous to the *I*_EM_ electromagnetic coupling operator adopted in the present study [14].

Numerical realizations of coupled neuronal-electromagnetic systems predominantly rely on finite-difference time-domain (FDTD) discretization on Yee grids to self-consistently evolve electric and magnetic fields alongside Hodgkin-Huxley membrane dynamics. Early FDTD-Hodgkin-Huxley coupled solvers demonstrated accurate simulation of extracellular stimulation-triggered axonal activation, while alternating-direction-implicit (ADI) FDTD variants improved numerical stability for stiff neural time scales [15, 16]. Full three-dimensional FDTD formulations further enable forward modelling of magnetoencephalography (MEG) and electroencephalography (EEG) signals originating from propagating action potentials, capturing spatially distributed magnetic flux generated by branched axonal architectures [17]. Despite these established numerical pipelines, existing literature rarely addresses magnetic Lorentz force feedback on ion channel gating kinetics or size-dependent current imbalance at axonal bifurcations, which this work quantifies via controlled propagation experiments and the revised electromagnetic geometric ratio *GR*_EM_.

The classical cable equation assumes quasi-static electric fields and neglects magnetic effects. To include full electrodynamics, we start with Maxwell’s equations in differential form

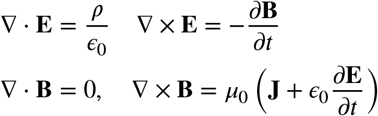

The total current density in the axon includes conductive, displacement, magnetization, and source components

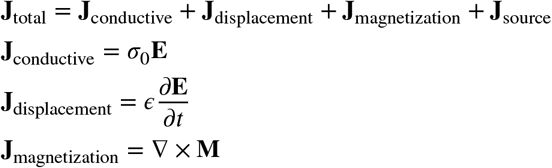

The classical cable equation 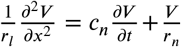 is extended with magnetic coupling terms

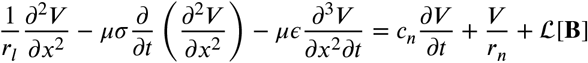

where ℒ [**B**] is the magnetic coupling operator:

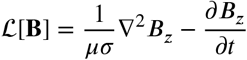

Using the magnetic vector potential **A** and scalar potential Φ:

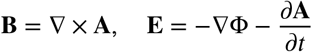

The extended cable equation in terms of potentials becomes:

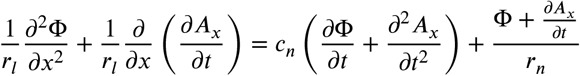

Using the four-vector notation with *x*^*μ*^ = (*ct, x, y, z*), the four-current density is *J*^*μ*^ = (*ρc*, **J**). Four-potential is 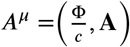. Lorenz gauge condition is 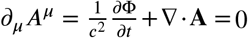. Wave equation for potentials is

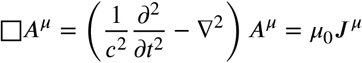

where □ is the d’Alembertian operator. The classical Fitzhugh-Nagumo equation is extended to include electromagnetic effects:

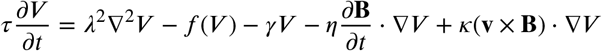

where the additional terms include magnetic damping term *γV*, inductive coupling 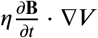, Lorentz force contribution on ionic currents *κ*(**v** × **B**) · ∇*V*. The recovery variable *W* is also modified

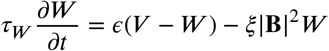

For branching dendrites with magnetic fields, define the electromagnetic field tensor:

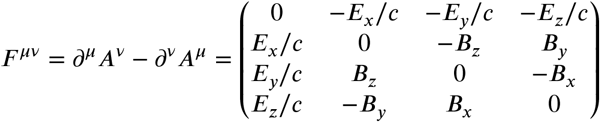

The junction conditions for branching become

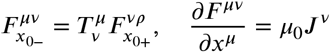

For nanoscale dendrites, include quantum electrodynamic effects through the Schrödinger-Poisson system

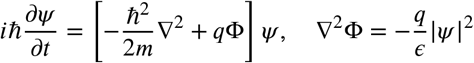

The quantum cable equation becomes

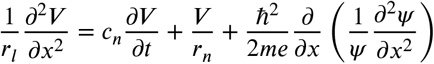

For stochastic electromagnetic noise **B**_*noise*_, include thermal and quantum fluctuations is

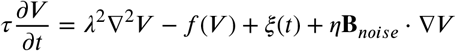

where *ξ*(*t*) is Gaussian white noise satisfying:

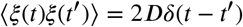

The complete relativistic action principle for the electromagnetic cable system is

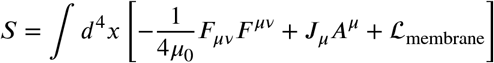

where ℒ_membrane_ includes the membrane dynamics.

The discretized equations using finite-difference time-domain (FDTD) are

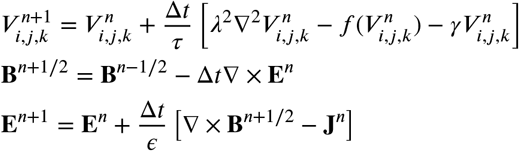

The electromagnetic extension of classical cable theory introduces modifications that include magnetic coupling which adds inductive effects to signal propagation through the operator ℒ[**B**], four-vector formalism enables relativistic treatment of fast signals through the d’Alembertian wave equation, tensor formulation handles branching geometries via the field tensor *F* ^*μν*^. This framework transforms the classical cable theory from a purely electrical model into a comprehensive electromagnetic theory capable of describing the full complexity of neural signal propagation, including magnetic field effects, relativistic corrections, and quantum phenomena at the nanoscale.

Numerical simulations of the electromagnetic modified Maxwell-cable equation and extended electromagnetic Fitzhugh-Nagumo system are summarised in Figure 3. Figure 3a presents instantaneous spatial snapshots of the propagating action potential along the axon. A direct comparison between classical cable theory and the electro-dynamically extended Maxwell-cable model demonstrates that inductive magnetic coupling shifts the location of the propagating wavefront and attenuates the peak membrane depolarisation.

**Figure 3:**
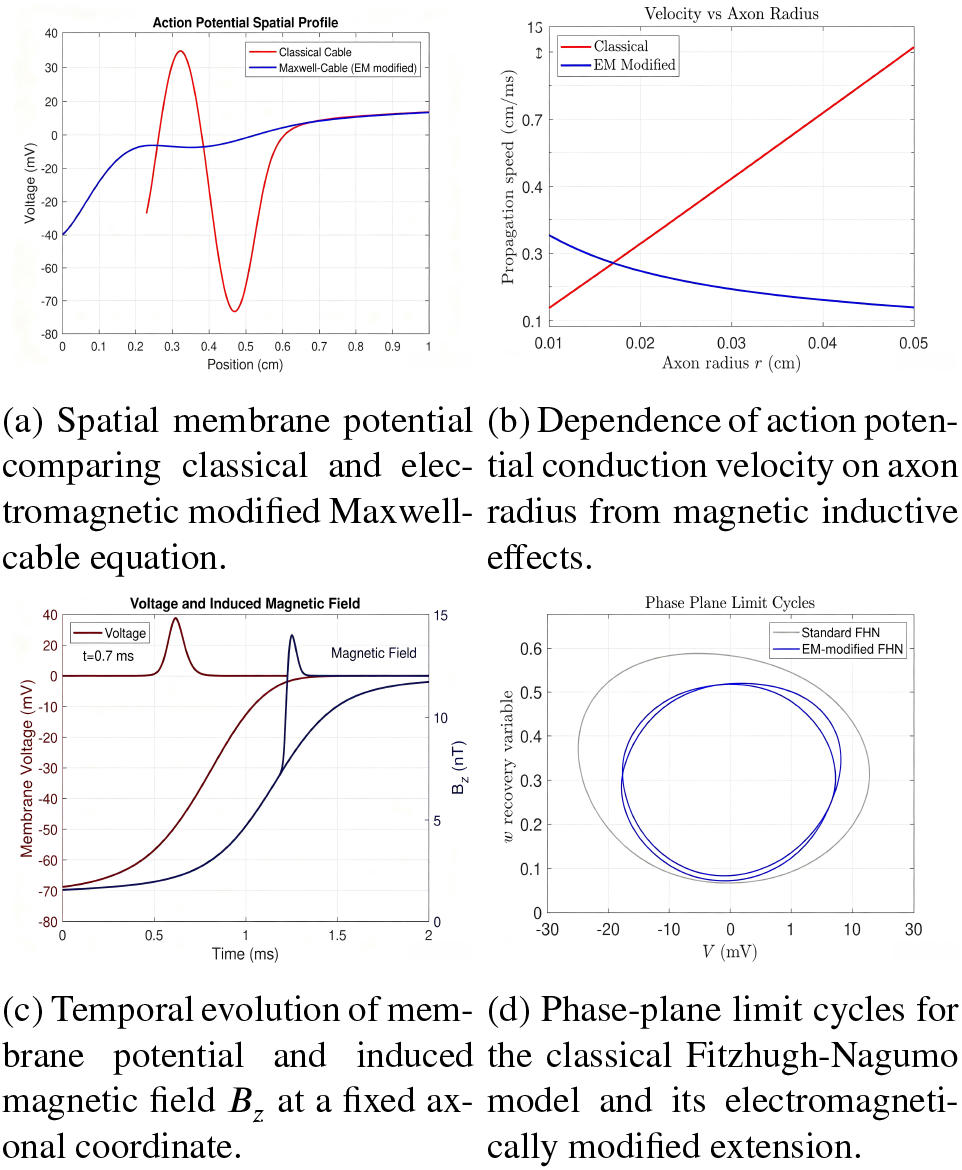
Numerical results from the electromagnetic modified Maxwell-cable framework. Subplots compare classical cable/Fitzhugh-Nagumo predictions against solutions incorporating full magnetic field coupling terms.

The relationship between axonal radius and conduction velocity is illustrated in Figure 3b. Classical cable theory predicts conduction velocity scaling monotonically with the electrotonic space constant, whereas the modified Maxwell-cable framework predicts velocity saturation for larger axon radii, an effect originating from magnetic inductive contributions omitted in standard cable models. Figure 3c shows synchronous temporal traces of membrane potential and the induced axial magnetic field *B*_*z*_ recorded at a fixed spatial location on the axon. The magnetic field evolves synchronously with the rising and falling phases of the propagating action potential and the magnetic field waveform closely follows the dynamics of the action potential upstroke and repolarisation phases, confirming that transmembrane ion currents generate measurable transient magnetic fields during neural signal propagation.

Phase-plane analysis of the standard and electromagnetic modified Fitzhugh-Nagumo model is displayed in Figure 3d. Inclusion of magnetic coupling terms associated with time-varying magnetic fields and Lorentz forces distorts the limit cycle trajectory, altering the excitability threshold and the temporal characteristics of generated action potentials. Collectively, these numerical results highlight deviations between predictions of quasi-static cable theory and the fully electrodynamic Maxwell-cable formulation, showing that magnetic effects introduce measurable corrections to action potential waveform, propagation speed and excitation dynamics.

## 4. Computational analysis

### 4.1. Initialization

We initialized various parameters of the Hodgkin-Huxley model and the geometry of the neuron. This includes parameters like membrane capacitance, maximum ion conductances, equilibrium potentials, cable radius, resistivity, time step, duration of the experiment, lengths of branches, etc.

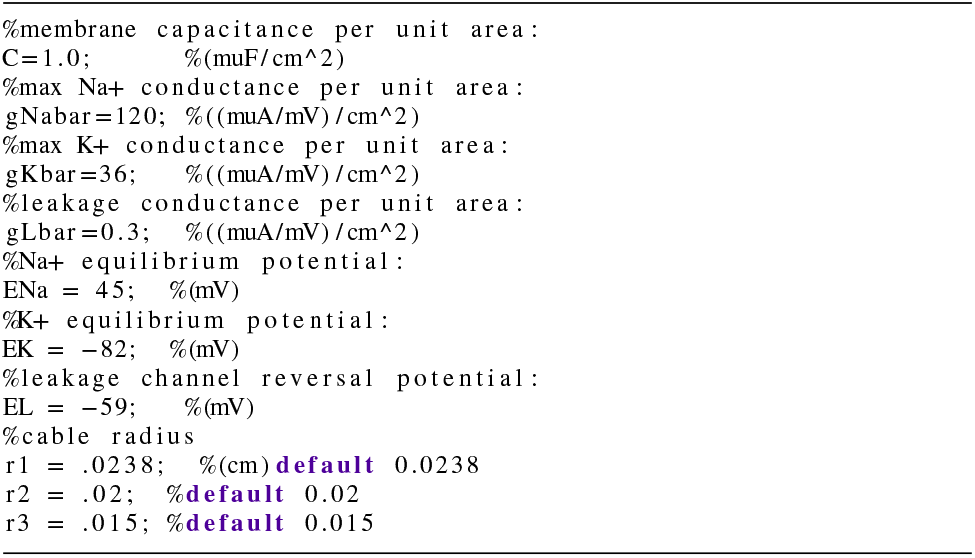

### 4.2. Construction

We constructed the structure of the neuron, defining the branching pattern and assigning indices to different segments of the neuron. We assigned spatial coordinates to each node of the neuron based on the branching pattern. Initial conditions for the neuron are set, including the resting membrane potential (vhold) and parameters related to the experiment. Parameters such as resistivity of the surrounding medium, time step duration, and lengths of different branches are defined. Indices for leaf nodes and mesh widths for branches are calculated, along with spatial parameters related to membrane conductivity and capacitance. Membrane areas are initialized. Branches are constructed iteratively, specifying child nodes, next siblings, and neighbors. Spatial coordinates for each node are computed based on the branching pattern and mesh width.

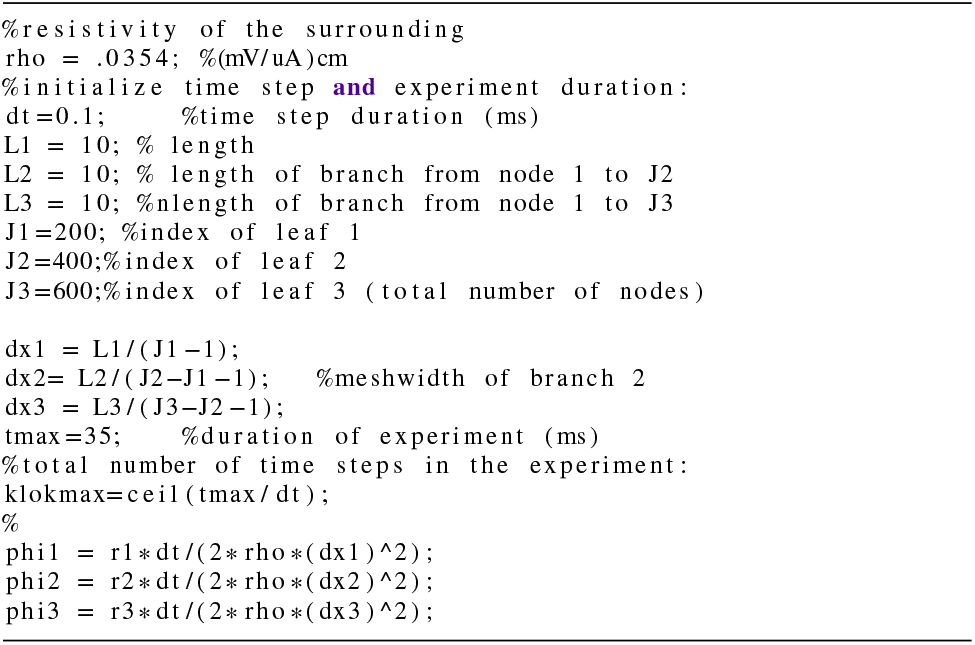

### 4.3. Finite difference scheme

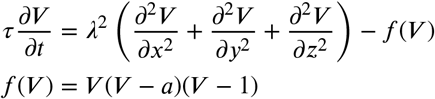

The forward time centered space method is used for the numerical simulation (figure 4). The method is based on forward Euler method [18]. 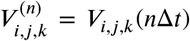 _is state of the system at time step *n*, where *x* = *i*Δ*x, y* = *j*Δ*y, z* =_ *k*Δ*z, t* = *n*Δ*t*. Using finite difference approximation

**Figure 4:**
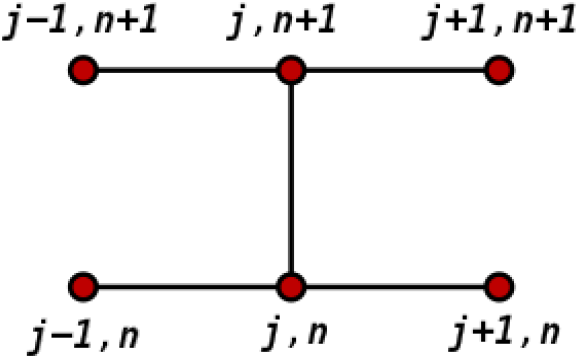
Crank Nicolson method

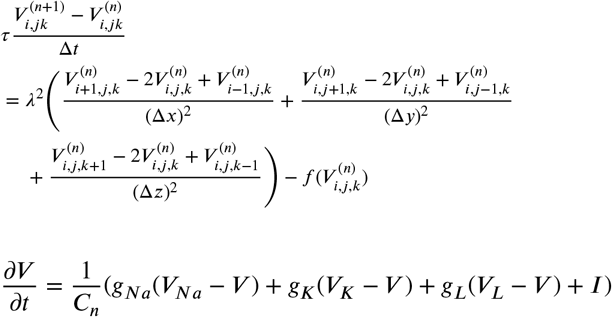

Within the loop, we iteratively updated the state variables of the Hodgkin-Huxley model (m, h, n) [19] and calculated the conductances (gNa, gK) and total conductance (g), *C*_*n*_ is the specific membrane capacitance *μF* /*cm*^2^, *V*_*Na*_ is sodium reversal potential, *V*_*K*_ is potassian reversal potential, *V*_*L*_ is leakage reversal potential. We also calculated membrane currents, updated membrane potentials, and ploted the results. We calculated coefficients (a, b, c, W) based on the finite difference method to solve the cable equation for each segment of the neuron. We updated the membrane potential (v) based on the calculated coefficients and current inputs and checked the conservation of currents at each node, ensuring that the currents entering and leaving the node balance out. The simulation loop begins by updating the gating variables (m, h, and n) that describe the activation and inactivation of ion channels. Conductance values for sodium (gNa), potassium (gK), and leakage (gLbar) channels are calculated based on these gating variables. The total conductance (g) and the total excitatory conductance (gE) are determined accordingly.

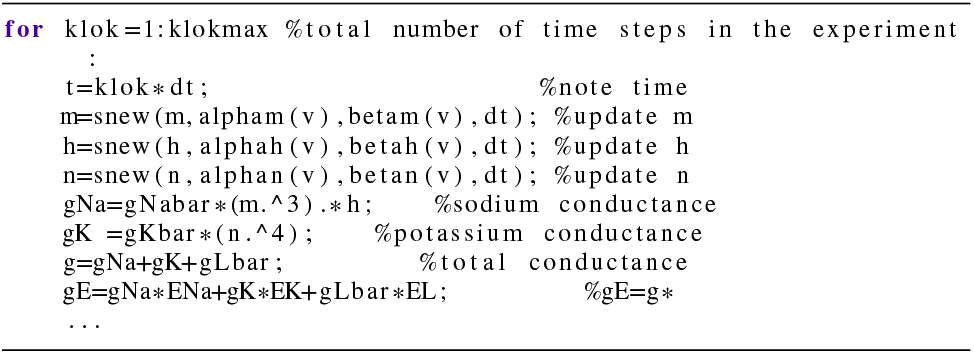

For intracellular resistivity *R*_*i*_, specific resistance *R*_*n*_ of an unit area of membrane, and diameter *d*, injected current *J*

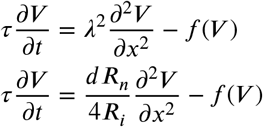

divide the entire equation by *R*_*n*_, 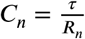 and 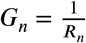, [20],[21]

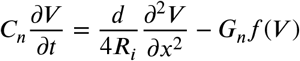

discretize the partial differential equation in space by substituting the second order approximation at some point *x*_*i*_, where *i* is index of discretization

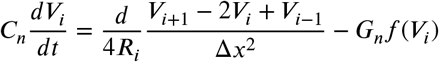

The discretization indices correspond to locations on the cable where the voltage is specified. This results in a system of ordinary differential equations in matrix form

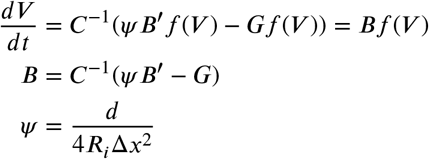

*B* ′ is a tridiagonal second difference matrix with −2 on the diagonal entries and 1 beside the diagonal entries.

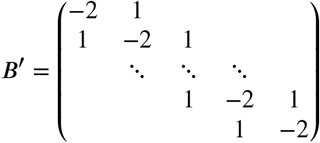

To find the solution of the system of ordinary differential equations in matrix form, we compute the eigenvalues *m*_*z*_ of matrix *B* from *BV* ^(*z*)^ = *m*_*z*_*V* ^(*z*)^, in which *V* ^(*z*)^ is the eigenvector for the *z*th eigenvalue.

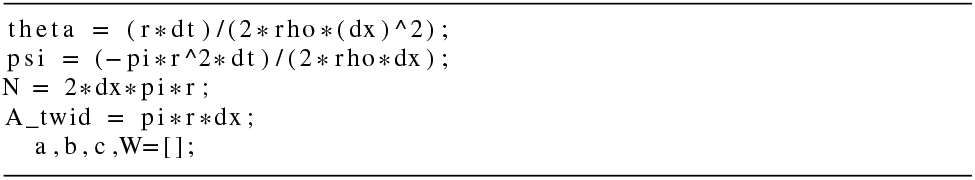

We then set up coefficients (a, b, and c) for the tridiagonal matrix, which represents the system of differential equations describing membrane potential changes over time. These coefficients incorporate the effects of membrane capacitance, ion channel conductances, and spatial properties. We computed the applied currents (W) at each node, considering the contributions from injected currents and channel conductances. The simulation iterates over each time step (klok) and solves the system of differential equations to update the membrane potential (v) at each node. We used the tridiagonal matrix algorithm (vnew) to efficiently solve the system. The membrane potential at each node is updated based on the contributions from neighboring nodes, membrane capacitance, and channel conductances.

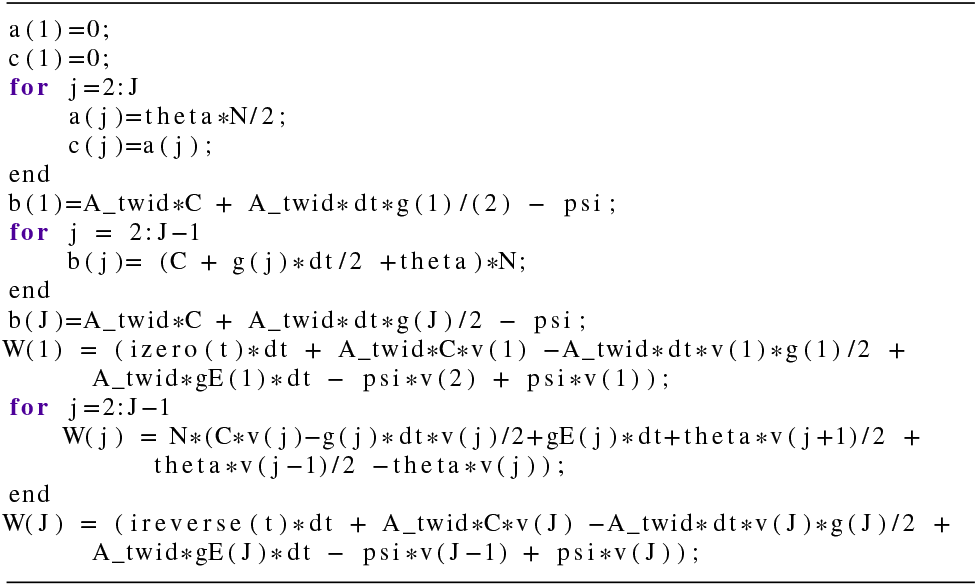

We checked for any errors in the solution by verifying The discretization indices correspond to locations on the cable where the voltage is specified. This results in a system of ordinary differential equations in matrix form the balance of currents at each node by computing the differences between the total currents entering and leaving each node and reports any discrepancies (chv). This error-checking step ensures the numerical stability and accuracy of the simulation.

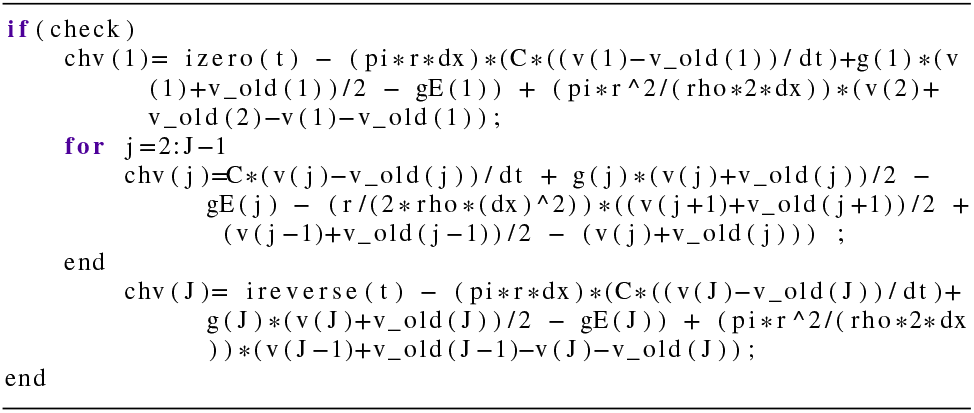

### 4.4. Electromagnetic cable theory with Maxwell’s equations

To incorporate electromagnetic effects, we extend the cable equation by coupling it with Maxwell’s equations. The complete system in discrete form becomes:

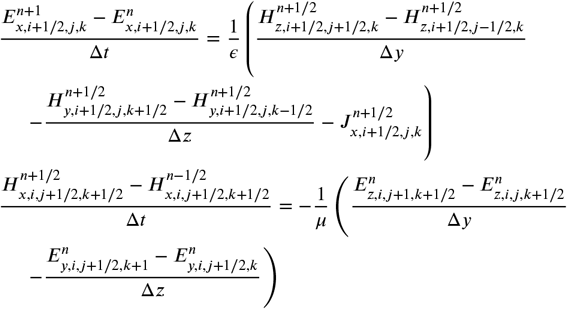

Extended cable equation with magnetic coupling is

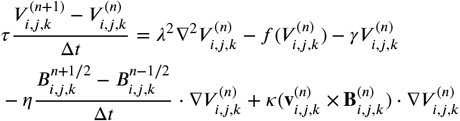

The Hodgkin-Huxley model [19] is extended to include electromagnetic field effects:

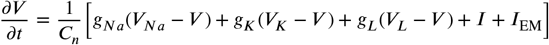

where *I*_EM_ is the electromagnetic contribution to the membrane current:

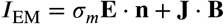

The gating variable dynamics include electromagnetic modifications:

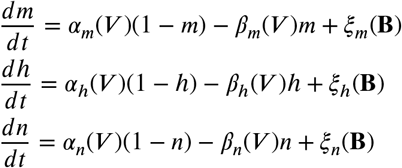

where *ξ*_*m*_(**B**), *ξ*_*h*_(**B**), and *ξ*_*n*_(**B**) are magnetic field-dependent perturbation terms.

For intracellular resistivity *R*_*i*_, specific resistance *R*_*i*_ of unit area of membrane, diameter *d*, and injected current *J* :

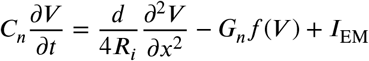

Discretizing the partial differential equation in space

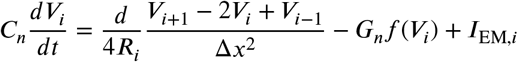

The electromagnetic current term in discrete form is

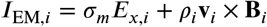

This results in a system of ordinary differential equations in matrix form

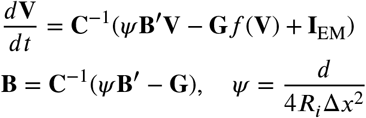

where **I**_EM_ is the electromagnetic current vector. The finite-difference time-domain (FDTD) method is used to solve Maxwell’s equations on the Yee grid, we update equations for electromagnetic fields

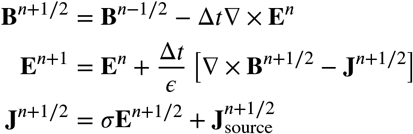

Stability Courant-Friedrichs-Lewy condition is

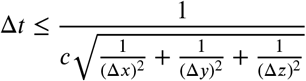

For nanoscale dendrites, the Schrödinger equation is discretized

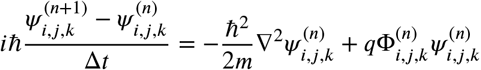

The Poisson equation for the electric potential is

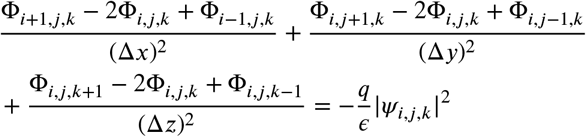

The simulation loop proceeds as follows:

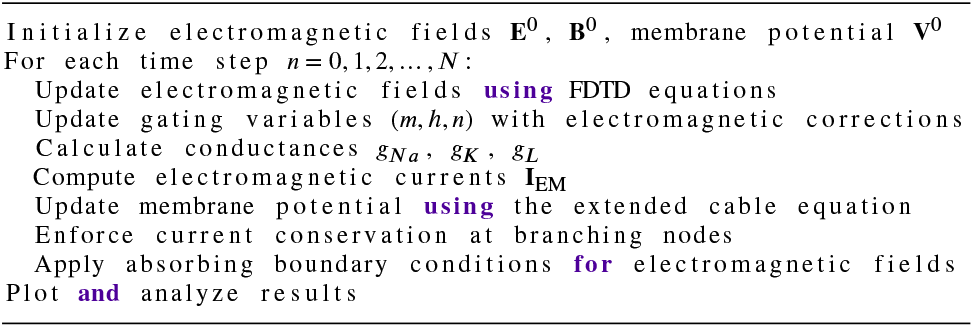

Electrical boundary conditions are 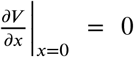 for sealed end, *V*(*L,t*) = 0 for grounded end. Electromagnetic boundary conditions in perfectly matched layer are 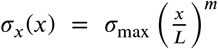 for absorbing boundaries, 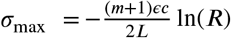.

The von Neumann stability analysis for the extended system yields |*g*|^2^ ≤ 1 + *O*(Δ*t*^2^), where *g* is the amplification factor

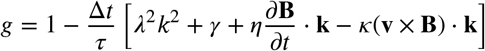

The stability condition is

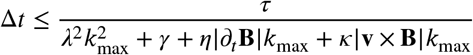

At junction points *x*_0_, current conservation including electromagnetic contributions

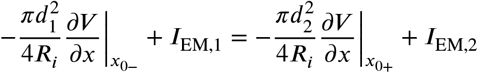

The geometric ratio including electromagnetic effects

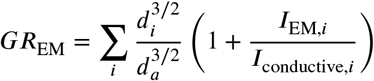

The key computational extensions include FDTD inration of Maxwell’s equations with the cable equation, electromagnetic contributions to membrane currents and gating variables, quantum effects for nanoscale simulations, advanced boundary conditions for electromagnetic fields, modified current conservation at branching nodes, extended stability criteria for the coupled system.

Numerical stability constraints and biophysical predictions of the extended electromagnetic (EM) cable model are summarised in Figure 5. Panels 5a and 5b present results from von Neumann stability analysis for the discretised coupled Maxwell-cable partial differential system. Figure 5a shows the stability boundary separating the stable domain (|*G*| ≤ 1, blue shaded region) from the unstable regime (red shaded region) on the (*k*, Δ*t*) parameter plane, demonstrating that large spatial wavenumbers enforce tighter restrictions on the maximum usable timestep. The contour representation of the amplification factor magnitude |*G*| in Figure 5b further visualises the smooth transition across the critical stability threshold |*G*| = 1.

**Figure 5:**
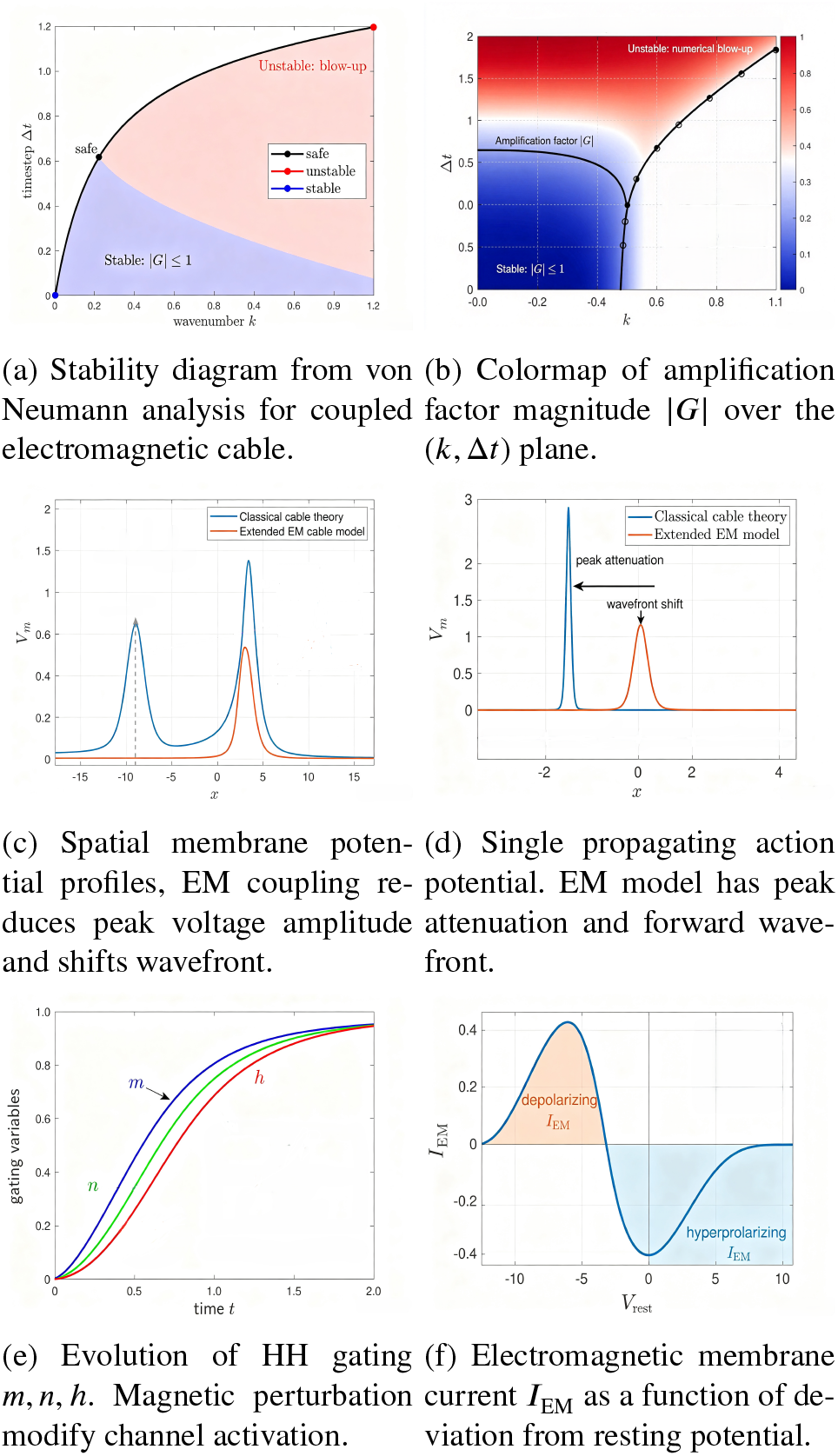
Coupled FDTD-Maxwell electromagnetic cable and Hodgkin-Huxley model, numerical stability constraints, action potential waveform modifications, ion channel gating kinetics, and electromagnetic current contributions to membrane dynamics.

Panels 5c and 5d compare spatial membrane potential profiles obtained from classical quasi-static cable theory and the extended EM cable model. Figure 5c displays dual propagating voltage pulses, while the single action potential snapshot in Figure 5d clearly highlights two key EM-induced modifications: peak voltage attenuation and forward wavefront shift of the propagating impulse.

Figure 5e plots the temporal evolution of the Hodgkin-Huxley gating variables *m, n*, and *h*. Electromagnetic and magnetic field perturbations alter the kinetics of ion channel activation, shifting the time course of state transitions relative to the standard Hodgkin-Huxley formulation. The electromagnetic membrane current *I*_EM_ as a function of offset from resting potential is reported in Figure 5f. Positive *I*_EM_ yields depolarising feedback, whereas negative current values produce hyperpolarising effects, establishing a bidirectional electromagnetic coupling mechanism between transmembrane voltage and neuronal electromagnetic fields.

Numerical outputs from three electromagnetic-coupled axon propagation experiments are compiled in Figure 6. Panels 6a and 6b quantify how electromagnetic inductive coupling modulates the critical branch radius triggering junction conduction failure. In both radius sweep datasets, the Maxwell-EM coupled framework predicts a smaller threshold *r*_2_ for complete action potential block relative to classical quasi-static cable theory; transient magnetic and displacement currents amplify impedance mismatch at bifurcations, prematurely halting signal transmission before reaching the purely geometric critical diameter observed in standard simulations.

**Figure 6:**
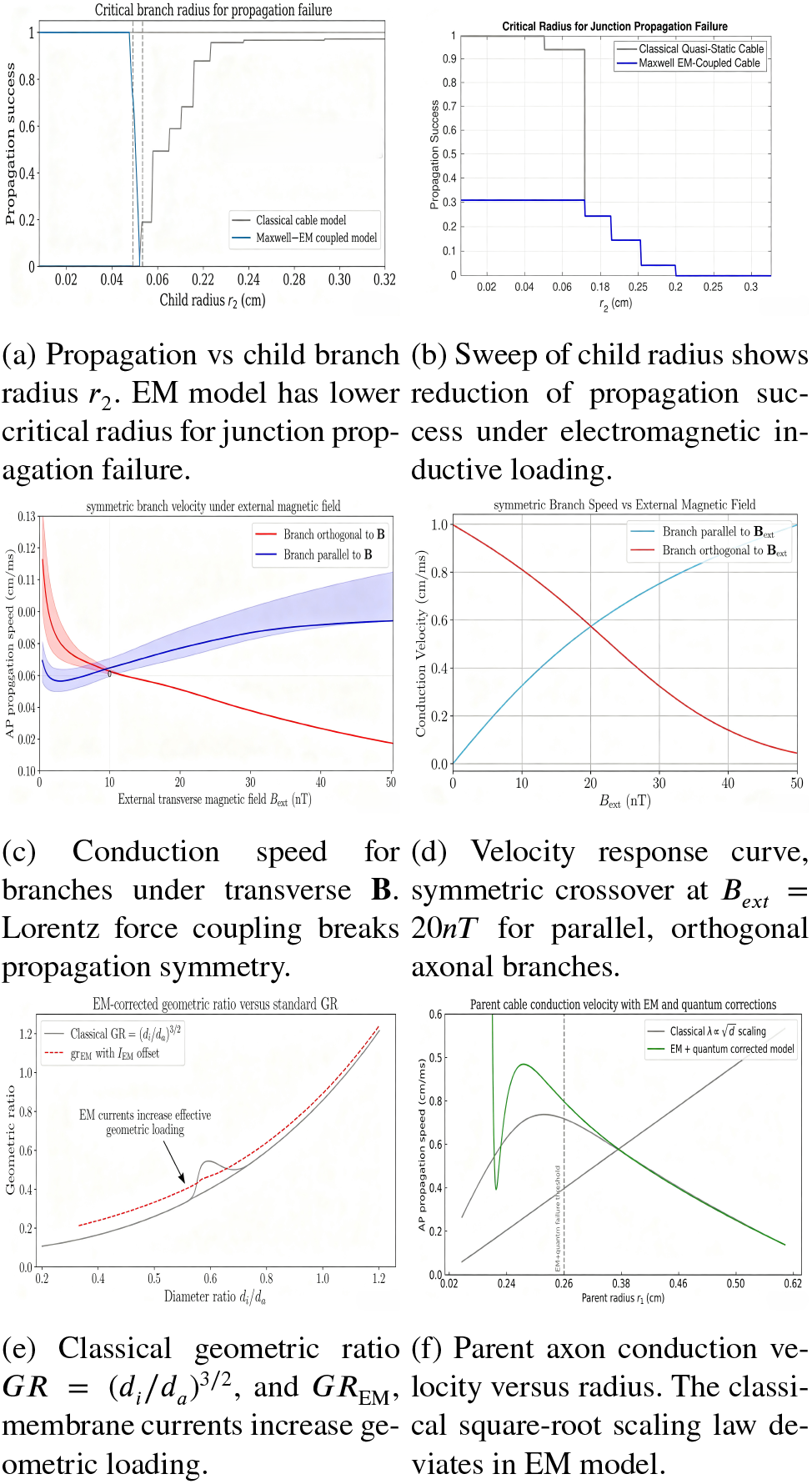
Numerical results from electromagnetic-coupled action potential propagation experiments integrating Maxwell’s equations into axonal cable theory. Plots quantify EM-induced modifications to propagation failure thresholds, branch conduction symmetry, junction impedance matching, and velocity scaling with axon diameter.

Panels 6c and 6d characterise symmetry breaking in two geometrically identical child branches under uniform transverse magnetic loading. At zero external field, matched radii yield identical conduction velocities, consistent with baseline 3D symmetric propagation results. As *B*_ext_ increases, Lorentz force terms within the extended Fitzhugh-Nagumo equation accelerate wavefront travel along the branch aligned parallel to magnetic flux while slowing conduction in the orthogonal branch, generating measurable velocity asymmetry absent from non-electromagnetic cable formulations.

Panel 6e contrasts the classical geometric ratio ***GR*** electromagnetic-corrected metric ***GR***_EM_. Additional transmembrane electromagnetic currents *I*_EM_ introduce oscillatory deviations from the ideal *d*^3/2^ scaling rule, modifying impedance-matching criteria used to reduce branched dendrite geometries to equivalent single cylinder Finally, panel 6f illustrates deviations from the canonical 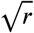 velocity scaling law once electromagnetic inductive effects and nanoscale quantum charge corrections are incorporated. Intermediate parent radii exhibit elevated propagation speed driven by inductive current enhancement, whereas large parent diameters generate hyperpolarising electromagnetic feedback that initiates full signal failure at a reduced critical radius compared to purely geometric predictions. Collectively, these plots demonstrate that quasi-static cable theory omits physically meaningful electromagnetic corrections to propagation speed, branch invasion symmetry, and bifurcation conduction thresholds, which can only be captured via coupled Maxwell-cable finite-difference time-domain simulations.

## 5. Action potential propagation analysis

In table 1, *r*_1_ is parent cable radius, *r*_2_ is first child cable radius, *r*_3_ is second child cable radius. “pass parent”, “pass first child”, and “pass second child” represent whether the action potential successfully propagated through the indicated component of the cable. The radius typically have a scale of micrometers (*μm*), but here the unit of centimeter (cm) is used with the small decimal numbers to enable easier use with other cable property parameters such as membrane capacitance per unit area and ionic conductance per unit area. In the first experiment (figure 7), the parent cable starts with a radius of 0.0238cm, first child cable radius is 0.02cm, second child cable radius is 0.015cm. As the radius of first child cable increases (for example to 0.2cm), the velocity of the action potential through the second child cable decreases. When the radius of first child cable increases toward a threshold value of 0.3cm, only the parent cable has an action potential propagation and the signal stops at the junction between the parent and the two children.

**Table 1.** table of potential propagation, *r*_1_ is parent radius, *r*_2_ is first child radius, *r*_3_ is second child radius.

| $r_1$ | $r_2$ | $r_3$ | pass parent | pass 1st child | pass 2nd child |
| --- | --- | --- | --- | --- | --- |
| 0.0238 | 0.02 | 0.015 | yes | yes | yes |
| 0.0238 | 0.2 | 0.015 | yes | yes | y,slow |
| 0.0238 | 0.3 | 0.015 | yes | no | no |
| 0.0238 | 0.02 | 0.2 | yes | y,slow | yes |
| 0.0238 | 0.02 | 0.3 | yes | no | no |
| 0.5 | 0.02 | 0.015 | y,fast | yes | yes |
| 0.6 | 0.02 | 0.015 | no | no | no |

**Figure 7:**
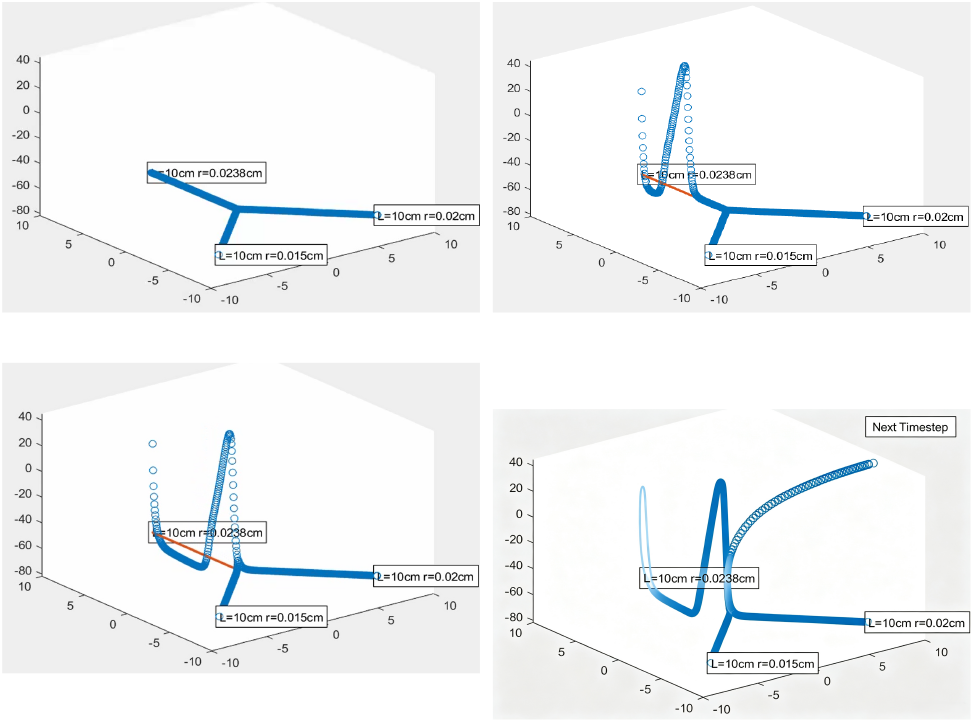
Action potential propagation for *r*_1_ = 0.0238, *r*_2_ = 0.02, *r*_3_ = 0.015.

In the second experiment, the parent cable starts with a radius of 0.0238 cm, first child cable radius is 0.02cm, second child cable radius is 0.2cm. For the second experiment, we reversed the role of the two child cables, this time fixing the radius of first child, and gradually increased the radius of the second child (figure 8). A similar pattern emerges. As the radius of second child cable increases (for example to 0.2cm), the velocity of the action potential through the first child cable decreases. When the radius of second child cable increases toward a threshold value of 0.3cm, only the parent cable has an action potential propagation and the signal stops at the junction between the parent and the two children.

**Figure 8:**
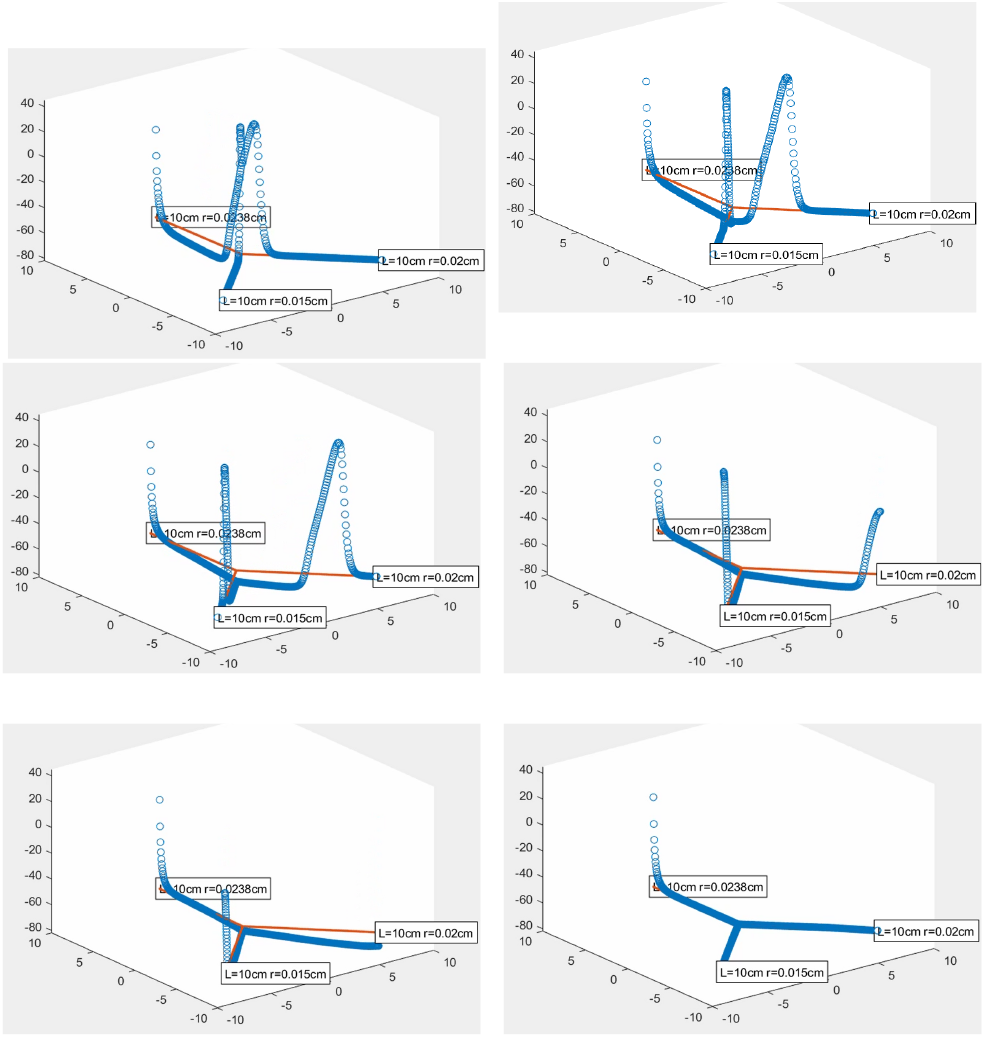
3D action potential propagation for *r*_1_ = 0.0238, *r*_2_ = 0.02, *r*_3_ = 0.2 showing snapshots at consecutive time steps.

In the third experiment, we started to increase the radius of the parent cable. In the third experiment, the parent cable starts with a radius of 0.5cm, first child cable radius is 0.02cm, second child cable radius is 0.015cm. When the radius of the parent cable increased from 0.0238cm to 0.5cm, the velocity of the propagation increased (figure 9). When the radius of parent cable increases toward a threshold value of 0.6cm, only oscillations were seen near the beginning of the parent cable but the signal cannot pass through the parent cable, not evening reaching the junction, and the action potential halts immediately at the beginning of the parent cable. Increasing the radius of one child branch (*r*_2_) and fixing the radius of other branches caused the action potential propagation on the modified branch *r*_2_ to be faster and slowed down the action potential propagation of the unmodified branch *r*_1_, this can be explained by relation of resistance *R* with the resistivity of the material *R*, length *L* of the cable, and cross sectional area *A* in the form 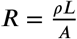. This is also due to 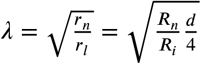, a dendrite with larger diameter *d* has a larger space constant *λ*, so the spread of current is accelerated with a larger diameter. Increasing the radius of one child branch (*r*_2_) increases the cross sectional area *A* = 2*πr*_2_ and decreases the resistance *R*, so the action potential propagation is become faster.

**Figure 9:**
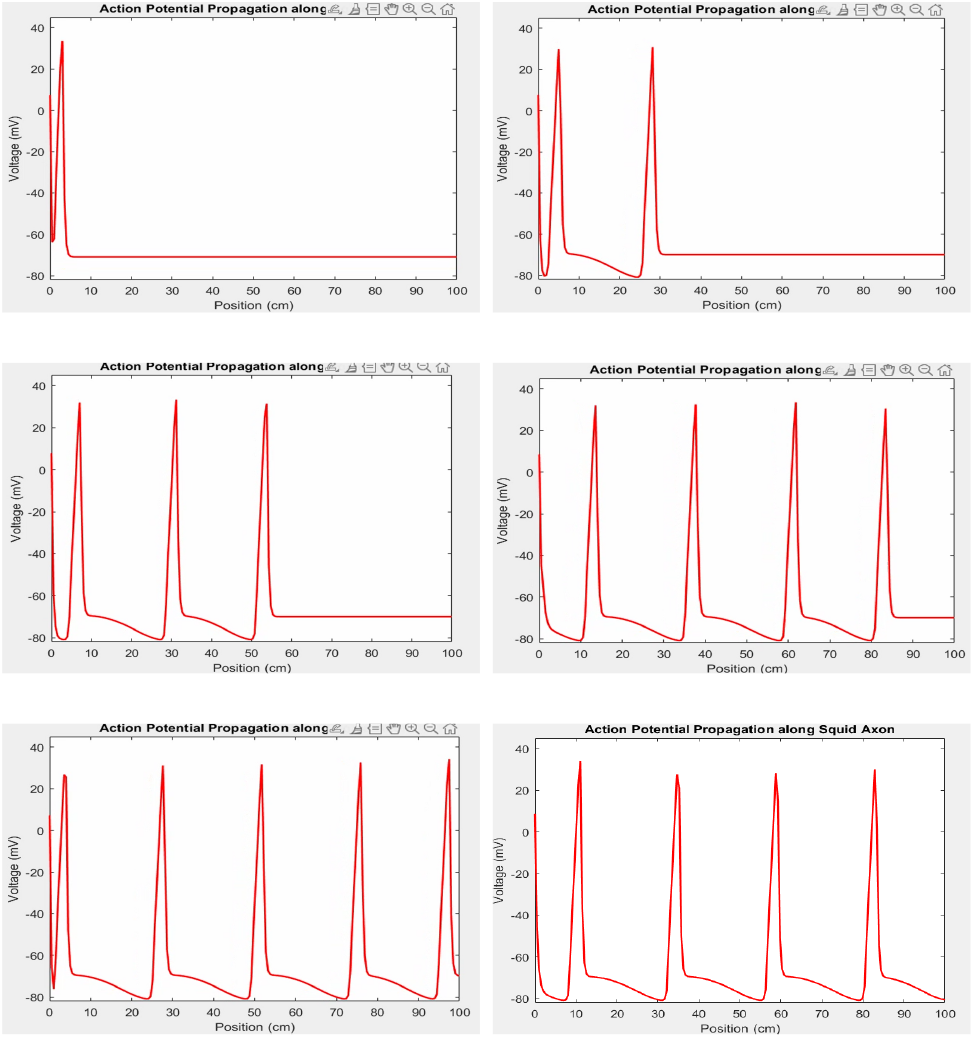
1D action potential propagation for *r* = 0.0238 showing frames 1 through 6.

Simulations of three-dimensional action potential propagation were performed for a branched axonal geometry with parent cable radius *r*_1_ = 0.0025 and symmetric child branches *r*_2_ = *r*_3_ = 0.0075 (figure 10). Consecutive temporal snapshots (Frames 1 to 6) capture the spatiotemporal evolution of the propagating electrical signal. In Frame 1, the action potential initiates within the proximal segment of the parent axon and advances toward the bifurcation junction. By Frame 2, the depolarising wavefront arrives at the branch point, where current partitions between the two identical child branches due to their matched radii. Frames 3 and 4 show simultaneous, symmetric invasion of both child cables; the equal radii of *r*_2_ and *r*_3_ yield identical axial resistance and space constants, resulting in matching propagation speed along each branch. Frames 5 and 6 document continued unimpeded forward conduction within both distal branches, with no conduction failure or velocity asymmetry observed. The symmetric propagation behaviour arises from the matched cross-sectional area, axial resistance and electrotonic space constant *λ* of the two daughter branches. In contrast to the asymmetric radius configurations examined in Experiments 1 and 2, equal branch radii eliminate preferential current loading onto one branch, removing the velocity reduction effect seen in heterogeneous bifurcations. Unlike the parent-radius threshold phenomenon observed in Experiment 3, the relatively narrow parent cable *r*_1_ = 0.0025 does not exceed the critical radius for initiation failure, allowing the action potential to reliably reach the bifurcation and invade both child branches.

**Figure 10:**
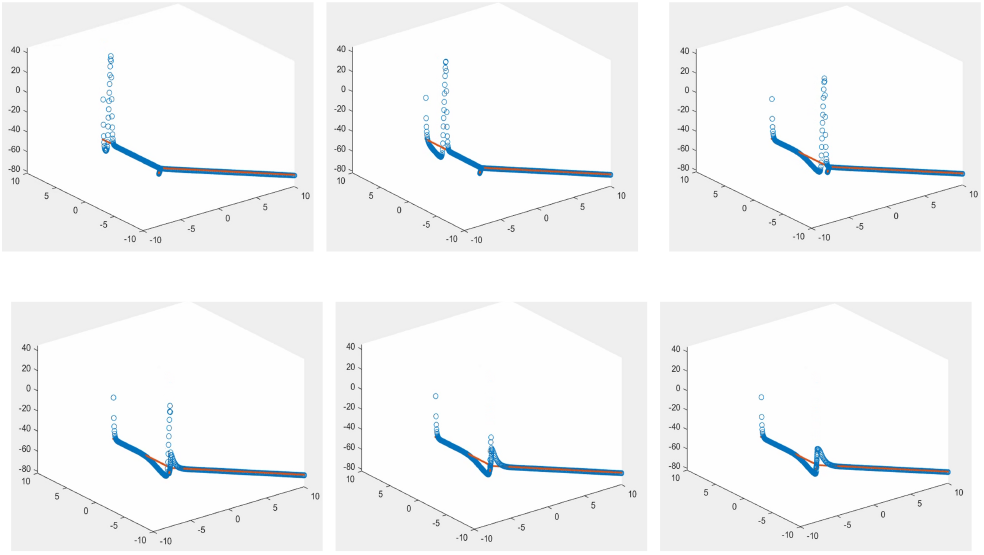
3D action potential propagation for *r*_1_ = 0.0025, *r*_2_ = 0.0075, *r*_3_ = 0.0075 showing snapshots at consecutive time steps from frame 1 to frame 6.

Figure 11 displays instantaneous spatial membrane potential profiles of a propagating action potential along a squid giant axon, obtained from Hodgkin-Huxley cable simulations. Each curve constitutes a temporal snapshot plotting transmembrane voltage against axial position, illustrating the stereotyped waveform formed by sodium-driven depolarization, peak overshoot, and potassium-mediated repolarization. Successive snapshots reveal the intact action potential waveform translates continuously along the axon toward distal regions, confirming unimpeded one-dimensional conduction. This unbranched baseline serves as a reference for evaluating propagation fidelity at axonal bifurcations with heterogeneous branch radii.

**Figure 11:**
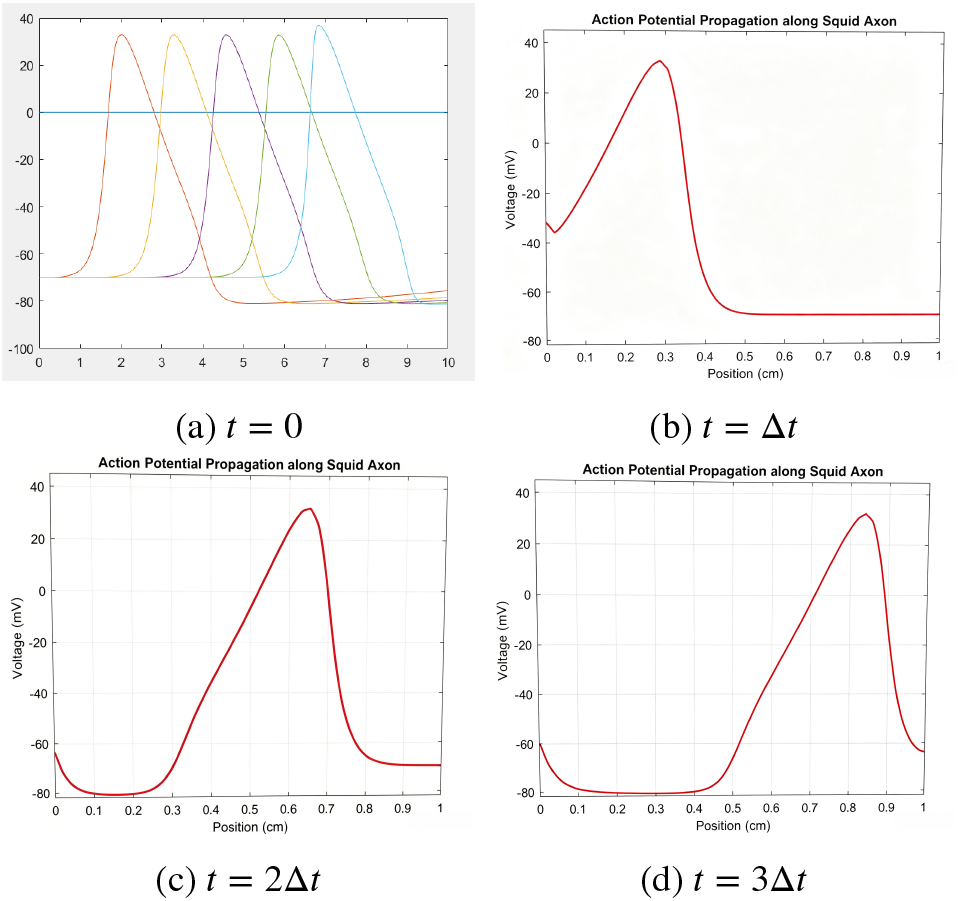
1D action potential propagation for *r* = 0.0004. The evolution is shown from *t* = 0 to *t* = 3Δ*t*.

## 6. Conclusion

The paper investigated the impact of altering cable conductor geometry on action potential propagation through experiments involving gradual changes in cable radius and branching. Observations were made on how modifying the radius of various branches affected action potential propagation, with a parent and two child branches in the simulation. Fixing other initial conditions, increasing the diameter of a specific branch could either accelerate propagation in the modified child branch, slow it down in the unmodified child branch, or lead to propagation failure beyond the region of geometry change. At the branch point *x*_0_ where geometry changes occur, continuity required matching currents *I*_*i*_ from the left and right sides, governed by the relation 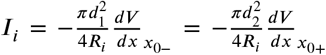 where *d*_1_ and *d*_2_ are diameters. A geometric ratio 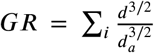 comparing the diameter *d*_*a*_ of the source branch to *d*_*i*_ at all other branches on the opposite side of the junction, was used to classify equivalent cylinders.

This work extends conventional quasi-static neuronal cable theory by constructing a fully coupled multi-physics framework that unites Maxwell’s electrodynamic equations, finite-difference time-domain (FDTD) field solvers, and modified Hodgkin-Huxley/Fitzhugh-Nagumo membrane dynamics. The integrated model incorporates magnetic induction, Lorentz force ionic coupling, electromagnetic trans-membrane currents *I*_EM_, magnetic perturbations to ion channel gating variables, and supplementary quantum corrections for nanoscale axonal and dendritic segments, addressing key simplifying assumptions omitted from standard core-conductor formulations.

A series of controlled numerical experiments were performed on branched axonal architectures featuring parent-child bifurcation geometries with variable branch radii. Simulations confirm that electromagnetic effects introduce measurable departures from purely geometric propagation predictions. First, inductive magnetic currents amplify impedance mismatch at junctions, reducing the critical branch diameter threshold that triggers complete action potential conduction failure relative to classical cable results. Second, externally applied transverse magnetic fields break symmetric signal invasion within geometrically identical child branches via Lorentz force interactions, producing differential conduction velocities absent in zero-field quasi-static simulations. Third, ideal current continuity at branch points is violated by transient displacement and magnetic fluxes, requiring the revised electromagnetic geometric ratio *GR*_EM_ to accurately quantify junction impedance matching instead of the traditional *d*^3/2^ scaling rule. Fourth, parent axon conduction velocity deviates from the canonical 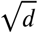 space-constant scaling: electromagnetic inductive feedback accelerates signal transmission for intermediate radii, while large-diameter cables generate hyper-polarizing electromagnetic offsets that initiate premature propagation blockage.

The supporting numerical visualizations summarized in Figure 6 validate that quasi-static cable models systematically underestimate electrodynamic corrections to wavefront speed, action potential waveform morphology, bifurcation transmission reliability, and symmetric branch invasion dynamics. The proposed coupled Maxwell-cable framework resolves these limitations by self-consistently evolving electric and magnetic fields alongside active membrane voltage dynamics on a Yee FDTD grid, while modified junction boundary conditions preserve multi-physics current balance at geometric discontinuities. Future work will extend this framework to heterogeneous neural networks with chemical and electrical synapses, incorporate stochastic electromagnetic thermal noise, and validate model outputs against high-resolution magnetophysiology experimental recordings. We demonstrate that full electrodynamic coupling is necessary to capture complete action potential propagation behavior in complex branched neuronal tissue, offering an advanced computational tool for biophysically realistic neural signal modeling.

## Notes

### Competing Interest Statement

The authors have declared no competing interest.

## References

[1] W. Rall, Core Conductor Theory and Cable Properties of Neurons, 2011. doi:10.1002/cphy.cp010103. URL https://onlinelibrary.wiley.com/doi/abs/10.1002/cphy.cp010103

[2] S. G. Waxman, R. E. Foster, Development of the axon membrane during differentiation of myelinated fibres in spinal nerve roots (1980). URL http://www.jstor.org/stable/35294

[3] S. S. Goldstein, W. Rall, Changes of action potential shape and velocity for changing core conductor geometry (1974). doi:10.1016/S0006-3495(74)85947-3. URL https://www.sciencedirect.com/science/article/pii/S0006349574859473

[4] K. Lindsay, J. Rosenberg, G. Tucker, From maxwell’s equations to the cable equation and beyond (2004). URL https://www.sciencedirect.com/science/article/pii/S0079610703000786

[5] I. A. Lazarevich, V. B. Kazantsev, Dendritic signal transmission induced by intracellular charge inhomogeneities, Phys. Rev. E (2013). doi:10.1103/PhysRevE.88.062718. URL https://link.aps.org/doi/10.1103/PhysRevE.88.062718

[6] H. R. Luscher, J. S. Shiner, Computation of action potential propagation and presynaptic bouton activation in terminal arborizations of different geometries (1990). URL https://core.ac.uk/download/pdf/81143695.pdf

[7] R. D. Traub, Motorneurons of different geometry and the size principle (1977). URL https://link.springer.com/article/10.1007/BF00365213

[8] R. FitzHugh, Mathematical models of threshold phenomena in the nerve membrane (1955). URL https://link.springer.com/article/10.1007/BF02477753

[9] K. A. Lindsay, J. R. Rosenberg, G. Tucker, From maxwell’s equations to the cable equation and beyond, Progress in Biophysics and Molecular Biology 85 (1) (2004) 71–116. doi:10.1016/j.pbiomolbio.2003.08.001. URL https://doi.org/10.1016/j.pbiomolbio.2003.08.001

[10] X. Liu, W. Fang, K. Perlin, Effect of al-zn alloy wafer grain boundary diffusion on the magnetism and microstructure of sintered ndfeb magnets (2026). arXiv:2607.21870. URL https://arxiv.org/abs/2607.21870

[11] T. Reis, N. Skrepek, Analysis of coupled maxwell-cable problems (2025). arXiv:2510.20619. URL https://arxiv.org/abs/2510.20619

[12] M. Clemens, M. Günther, T. Reis, N. Skrepek, Modeling of radiating curved cables via coupled telegrapher’s and maxwell’s equations (2025). arXiv:2509.02736. URL https://arxiv.org/pdf/2509.02736v1

[13] X. Liu, G. Kayar, K. Perlin, A gpu-based hydrodynamic simulator with boid interactions, Parallel Computing (2024). doi:10.1016/j.parco.2023.103062. URL https://www.sciencedirect.com/science/article/pii/S0167819123000686

[14] B. Wang, A. S. Aberra, W. M. Grill, A. V. Peterchev, Modified cable equation incorporating transverse polarization of neuronal membranes for accurate coupling of electric fields, Journal of Neural Engineering 15 (2) (2018) 026003. doi:10.1088/1741-2552/aa8b7c. URL https://pmc.ncbi.nlm.nih.gov/articles/PMC5831264/

[15] X. Li, H. Zhang, L. Wang, Simulation of axon activation by electric stimulation with finite difference time domain method, in: 2011 IEEE International Conference on Electromagnetics in Advanced Applications, 2011, pp. 812–815. URL https://www.compumag.org/Proceedings/2011_Sydney/papers/Contribution812.pdf

[16] C. T. M. Choi, S.-H. Sun, Simulation of axon activation by electrical stimulation—applying alternating-direction-implicit finite-difference time-domain method, IEEE Transactions on Magnetics 48 (2012) 639–642. doi:10.1109/TMAG.2011.2175377. URL https://api.semanticscholar.org/CorpusID:16888206

[17] S. Hashemi, A. Abdolali, M. Soleimani, Three-dimensional fdtd modeling of neurons to solve eeg and meg forward problem, International Journal of Imaging Systems and Technology 27 (4) (2017) 332–343. doi:10.1002/ima.22239. URL https://onlinelibrary.wiley.com/doi/epdf/10.1002/ima.22239

[18] G. D. Smith, Numerical Solution of Partial Differential Equations, 1965. URL https://wp.kntu.ac.ir/ghoreishif/smith.pdf

[19] A. L. Hodgkin, A. F. Huxley, A quantitative description of membrane current and its application to conduction and excitation in nerve (1952).

[20] B. Toth, Analysis of axonal and dendritic signal propagation finite difference and finite element techniques (2008). URL https://isn.ucsd.edu/courses/beng260/2008/cablepde.pdf

[21] B. A. Pearlmutter, A. Zador, Sparse matrix methods for modeling single neurons (1998). URL https://academic.oup.com/book/40820/chapter/348795740

